# A Modular Photosynthetic Microbial Fuel Cell With Interchangeable Algae Solar Compartments

**DOI:** 10.1101/166793

**Authors:** Daniel Fleury

## Abstract

This project trial provides a novel small-scale solar harnessing technology which increases environmental effectiveness while maintaining optimal energy efficiency. Although modern solar panels are purposed in producing clean energy, the materials and byproducts of solar cell manufacturing are not eco-friendly. Thus, considering an organic, renewable and energy efficient solar cell model is necessary. Investigations explored multiple highly-photosynthetic algal species which were later integrated into a controlled microbial fuel cell system (MFC). The MFC1 contained algae culture species, including Chlorella Vulgaris, Nannochloropsis, and Spirulina. Parameters, such as periodic lipid yields, algal biomass, and light absorption were assessed throughout the cultivation process while maintaining a controlled environment. After 30 days of cultivation, the culture was transferred to an anode chamber in a closed loop small-scale MFC. Following the first day of algae transfer, microwatt output was analyzed from independent test trials. Statistical comparisons were drawn between electrical energy and light absorption, finding a generally positive correlation. Thus, it is concluded that mid-high algae concentrations significantly increased electrical micro-wattage in highly absorptive algal cultures. The optimal electric levels occurred at 286 A (absorbance) and 35 mW-Nannochloropsis, 123 A-Chlorella (30.2 mW), and 142 A (31 mW)-Spirulina culture. Due to higher absorption rates in the Nannochloropsis culture, this corresponds with the record high voltage levels. The analysis of data indicates that Algae-based MFCs are proven hopeful for alternative high-yield energy production.

## INTRODUCTION

The international standard of the energy sector has been concerning in its immediate and long term environmental impact. More specifically, the continuing environmental and ecological effects of mediums such as fossil fuels and coal output have influenced phenomenon such as global warming and the greenhouse effect. However, private and public energy corporations have experienced slow transitions into renewable energy. The undeniable root of this lagging changeover comes from world energy demands in growing populations. Although the most renewable techniques of energy delivery have been hydropower and biomass, these methods are unable to sustain mass populations due to its small scale output systems. The economical magnitude of biomass and hydropower is costly, due to its geographical limitations (i.e. costly land expansion). Furthermore, due to the finite availability of biomass, satisfying full energy demands alone is not possible. Additionally, when considering small scale biomass energy generation, net energy is easily lost due to the system’s high energy needs. Moreover, the combustion of extraneous or usable biomass can harm the environment by discharging carbon dioxide, encouraging possible climatic change. As a result, energy sustainability, consistency, and environmental efficiency, are not fulfilled. Similarly, the high yielding biofuel industry poses a plethora of economic drawbacks. The biofuel sector presents a demanding cost of production in contrast with the fossil fuel industry. The ecological facet of biofuel and biomass extraction is also penetrated by the ineffectiveness of monoculture and the use of fertilizers. Furthermore, systematic monoculture may deprive soiled nutrients by cultivating a continuous crop species/genus. In other words, the variety of crop species is bottlenecked, which reduces nutrients levels. Extraneous fertilizers may also be deleteriously discharged during the cultivation process, damaging the surrounding ecosystem. Most importantly, considerable net energy gains cannot be absorbed with small scale structures and frameworks of biofuel production. Constructing a compact and energy efficient utility (in general) for individuals is impractical with the previously acknowledged energy sources (coal, fossil fuel, biofuel/biomass). Although industrial scale energy is essential, a scalable energy system is crucial in all frames of worldwide economic demand. The current canvas of energy production presents insufficiency with energy scalability due to the lack of adjustability and interconnection. The current systems of energy production do not provide modulation nor convenience in partitioning energy into separate units.

## BACKGROUND AND REVIEW OF LITERATURE

Energy is a crucial resource proportional to the growth of individual populations around the globe. The high demands of exponentially growing groups and populations have constrained the options of energy production. More specifically, industrializing energy became essential in meeting population needs and demands. Economically, both fossil fuel and coal production are the most sufficient fields of immediate energy availability and output. According to a 2015 energy consumption report from Blue Sky Energy Partners, the combination of petroleum and coal expend nearly 52% of all energy demands in the United States-36% Oil and 18% Coal (U.S. Energy Information Administration, 2016). In addition to this national survey of energy consumption, a global review found nearly 62% of all needs accounted for coal and oil-33% Oil and 29.2% Coal (Columbia University, 2016). International and National reviews of energy consumption indicate that energy corporations are ultimately straying away from renewable energy types. The frame of renewable energy is hindered technologically, as it is limited by surrounding environmental conditions. More specifically, technologies, such as solar cells/solar panels, are constrained by the sunlight exposure in specific regions. The limitations of the renewable and nonrenewable energy sector create economical barriers, oftentimes being excluded from developing (3rd world) countries. Studies have presented a close correlation between both economic growth and energy availability (Neto et al., 2014). Due to the economic circumstances of 3rd and 2nd world countries, continuous access to both nonrenewable and renewable energies is constricted, due to financial limitations of large capacity/quantity. Contrary to non-renewable energy types, renewable sources do not suffice the energy market and the economy due to high initial costs and inconsistency (Giraldo et al., 2014). The economic and technological impediments of large-scale and non-intuitive renewable energy sources can be most magnified in impoverished and developing nations of Africa. Kenya is a developing country which necessitates biomass and waste combustion as the prime source of fuel. However, individual houses and facilities experience frequent power outages due to irregular Electricity production. The central drawback of irregularity lies in large-scale energy inconsistency. In other words, developing countries are oftentimes deprived of a smart grid system which provides stable energy flow to individual facilities, departments, and residents. Implanting a Smart Grid2, along with Distributed Generation3, may provide dependable energy flowage. More specifically, Distributed Generation may integrate electricity and power with small-scale technologies, which lessens stress on the central grid system (Vandaele & Porter, 2015).

## BIOLOGICAL SOLAR CELLS

Biological solar cells have been of particular research interest, due to the integration of metabolism and its implications in energy production. This system amalgamates a single species of live microbial organisms, such as anaerobic bacteria. However, biological materials such as Eukaryotic organisms (plantae) can be exploited through their photosynthetic and respiratory properties. Most importantly, the metabolic pathways can be manipulated by exploiting extraneously discharged electrons and protons in the media. As an alternative energy source, Microbial Fuel Cells advantages the sectors of energy and environmental sustainability by catalyzing naturally produced energy in growable living conditions (e.g. bioreactor and photobioreactor). Traditionally, the system houses three divisions/containments-An isolated Anode chamber, an isolated cathode chamber, and a semipermeable proton exchange membrane. In addition, the two electrodes (anode and cathode) are interconnected with conductive wires to form a fully attached simple circuit. The circuitry system is then completed by the particle transfer of the proton exchange membrane (PEM). Combining these three components forms a bioelectrochemical system. As bacteria is submerged into the anode chamber, freely electrons penetrate through the wall of the semipermeable membrane into the cathode containment, outputting electrical current. The proton exchange membrane constantly transports hydrogen ions to harness electricity. However, due to the vulnerability of these biological sources (regular pond bacteria) it is prone to contamination. Therefore, achieving a much more stable and adaptable microorganism is necessary. In addition, the metabolic processes of natural bacteria (mostly found in pond ecosystems) have varying metabolic processes. Thus, seeking a microorganism with great photosynthetic processes and cell respiration is necessary.

## RATIONALE

Investigating the photosynthetic nature of a novel algae-based biological solar cells as an alternative to currently manufactured solar panels is crucial. However, the current lack of environmental efficiency must be resolved in the modern solar cell model. As a most likely candidate, microbial fuel cells serve as source of novel experimentation and test sampling. Microbial fuel cells, as indicated in recent test trials, harnesses the energy by exchanging Hydrogen Ions from cellular respiration. However, the significance of photosynthetic algal cultures, containing Nannochloropsis, Spirulina, and Chlorella, will be investigated as a substitute for traditional bacteria. The environmental effectiveness of MFCs creates widespread economic potential in energy production and absorption through the sustainability of small and large scale electrical demands. A regenerable solar cell with the use of multiple algal species will simultaneously deplete carbon dioxide and produce high lipid yields/high electrical output from an MFC system.

However, situations such as the regenerability, cost effectiveness, and renewability are still prevalent. By manipulating the operations of a microbial fuel cell, energy output may be maximized by mimicking bioreactor culture conditions.

Algae can proliferate at high frequencies, which makes regenerability a big factor: The regenerability of this microorganism will eventually accumulate biomass which can be exploited for separate purposes such as agriculture (agricultural services can use biomass for cultivation). The photobioreactor component of this experiment will include a controlled environment. In addition, by using a photobioreactor, this ensures that there is a harnessing of solar-based energy. Exploiting biological materials is the criteria of this project due to the considerable availability of media in varying ecosystems. The high magnitude of Hydrogen Ion discharge may advantage the energy and environmental sectors of a small-large scale economy through easy configuration and clean energy access.

## METHODS AND PROCEDURES

The initial protocol was first initiated with containment of Algae culture in the Microbial Fuel Cell structure. A Microbial Fuel Cell with a suitable exchange of Hydrogen Protons and Electrons across the Proton Exchange Membrane (PEM) was crucial to fully harness the electrochemical properties of each Algal culture-Chlorella Vulgaris, Spirulina, and Nannochloropsis. Constant variables were maintained by precisely quantifying internal conditions and additional attachments (the PEM). The most basic constants (remaining throughout complete experimentation) included anodic and cathodic temperatures. Additionally, the concentrations of Algae culture were regulated at 1 mL of Algal Culture/10 mL of media. The Algal culture of each genus was obtained at high concentrations of 8 mL/10 mL of initial media. More specifically, additional nutrients media was added at concentrations of 1 mg/250 mL. The most critical constant variables and parameters are listed below, with sufficient range levels to correctly maintain culture growth:

**Table 1.0:** Essential (constant) parameters and conditions regulated in the Algal MFC cultures. Parameters are presented with the most optimal ranges/conditions for cultivation. All Algal cultures were continuously maintained within these minimum to maximum ranges.

| Parameter Regulated | Optimal Range |
| --- | --- |
| Temperature (°C) | 16-27 |
| Salinity (g.l-1) | 12-40 |
| Light intensity (lux) | 1,500-10,000 |
| Photoperiod (light: dark) Ratio in Hours | 16:8 - 24:0 (minima - maxima) |
| pH | 8.2-8.7 |

The most essential procedures included MFC construction, Algae cultivation and maintenance, Electrical testing, and supplementary Oil Yields (optional to experimentation). MFC construction involved attachable/interconnected 450 mL Anode and Cathode containers to fulfill the needs of the Algal culture-Cathode interactions. The Proton Exchange membrane was further composed of an Agar solution brought to boiling point and later froze at 0°C (32°F). The Cathodic chamber was comprised of an isolated and sterilized (distilled) continuously aerated water. Following the configuration of the Anodic and Cathodic chamber, with the integration of the PEM, MFC underwent electrical testing with a traditional multimeter to detect variations in output levels in each culture chamber. More importantly, direct correlations between Algal concentration and electrical levels (mW) were analyzed and discussed.

### MFC Construction: Designing the Anode and Cathode Chambers

The most simplified design of the MFC containment system involved interlinking both the Anode and Cathode chamber, sandwiched with a Proton Exchange Membrane. The parameters of both chambers were equalized to maintain controlled conditions and environments in each container. Both chambers held a capacity of 450 mL of Distilled water (Cathode) and Highly Concentrated Algal culture (Anode). In preparation for the PEM, ~10.16 cm diameter circular openings were made, consistent with the shaping of the circular PEM. The designing of the PEM system will be further discussed in its specified procedure. The “suctioning” nature of the Proton Exchange Membrane enabled both the Anode and Cathode chamber to converge without any major or minor leakages to interfere with the exchange process. Conveniently, the purchased tupperware containers were attachable, making reconfiguration and modification of the system possible. This further enhanced the theme of interchangeability and attachability, making immediate immediate configurations to the energy system possible. After nearly 50 mL of highly concentrated algae culture was obtained, the culture was immediately transferred to a 500 mL nutrient rich bioreactor medium composed of an initial 2 mg of All-Purpose growth media (1 mg/250ml). Parameters, such as cell concentration, pH, aquatic temperature, salinity, sterilization, and lux levels were assessed. Cell concentrations were obtained with a spectrophotometer. After two days of cultivation, the algae was transferred to a 450 mL MFC anode chamber. Electrical testing was executed with a multimeter.

The transfer system of the Microbial Fuel Cell ensured that the isolation of cultures was maintained. Furthermore, the containment of any possible contaminants was prevented with the architectural nature of the microbial Fuel Cell. Each chamber of the MFC was interconnected with pvc piping network, regulating and aggregating the nutrient and biological media of the Anodic and Cathodic regions. Additionally, the input and output exchange system of the fuel cell made configuration, attachability, and addition/reduction possible. In other words, adjusting critical parameters and concentrations was straightforward through the convenient architectural properties of the MFC. The Anodic and Cathodic chambers were constructed to mimic the regulatory conditions of a bioreactor. More specifically, the Anode chamber ensured stabilization of parameters and critical variables in culture and media to ensure controlled chamber conditions.

### PEM Construction: Designing an Electrolytic Passageway in the Microbial Fuel Cell

The current protocol for PEM construction comprises of a mixture of electrolytic polymers which facilitate the exchange of protons across a semipermeable membrane. The nature of the PEM satisfies the electrical exchange (Hydrogen proton transfer) between the Anode and Cathode chambers. More specifically, the PEM harnesses and manipulates freely moving protons (primarily in the Anode chamber) to output electrical energy from the bridgeway of the two submerged electrodes in the Anode and Cathode chambers. The electrolyte membrane gradient between the Anodic and Cathodic chambers optimized efficiency and electrical production by acting as a continuous bridgeway of Hydrogen Proton exchange. The medium of the Proton Exchange Membrane is comprised of a simplified Agar-based solution, including a mid-high concentration of sodium. The reactants of the PEM were freely adjusted in association with the experimental trials. In other words, based on the conditions of Algal culture, PEM consistency and concentrations were regulated for optimization. The following table describes the mixture of the PEM medium mixture.

**Figure 1:**
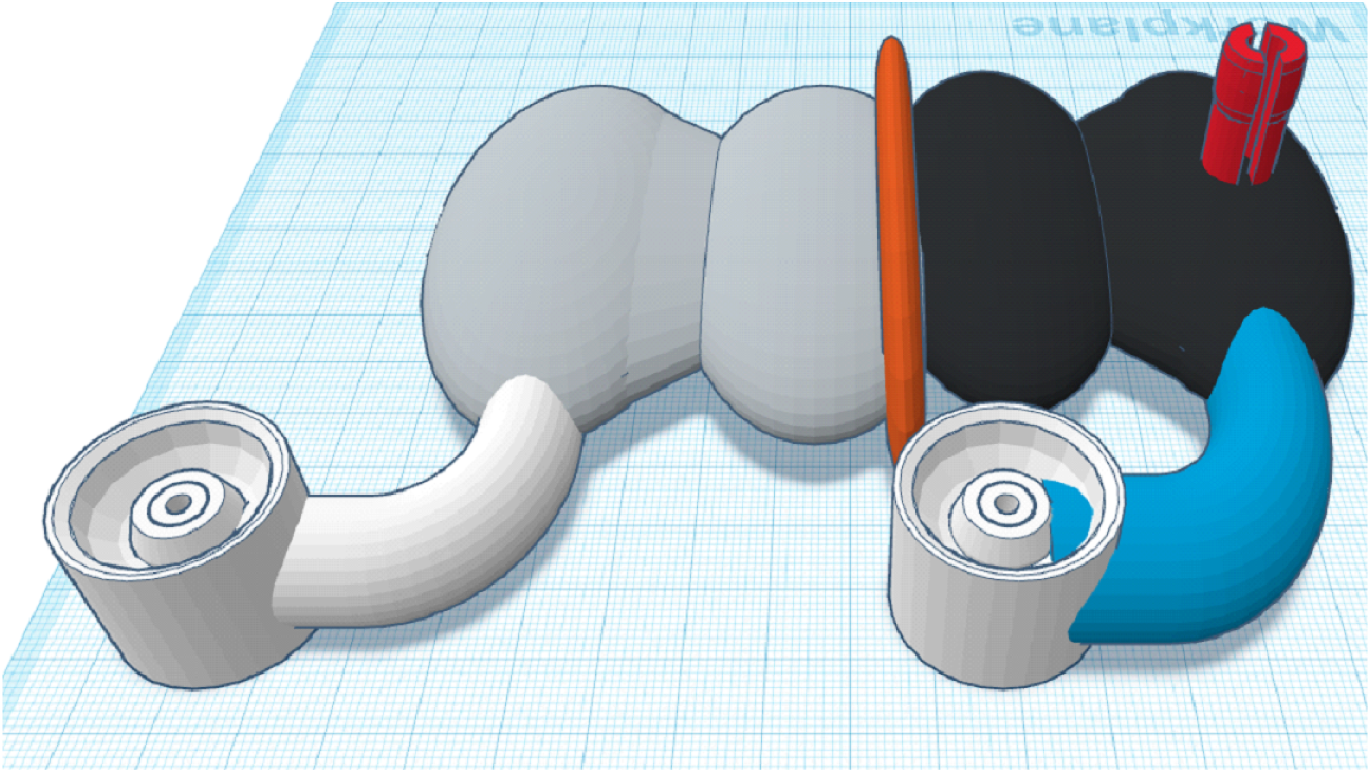
The image presents a 3D rendering of the construction of the interlinked Microbial Fuel Cell system. Additionally, attached tubing, the Proton Exchange Membrane, and O2 exhaust is exhibited. The Cathode chamber is featured on the left of the diagram, whereas the Anode chamber is displayed on the right of the diagram.

**Figure 2:**
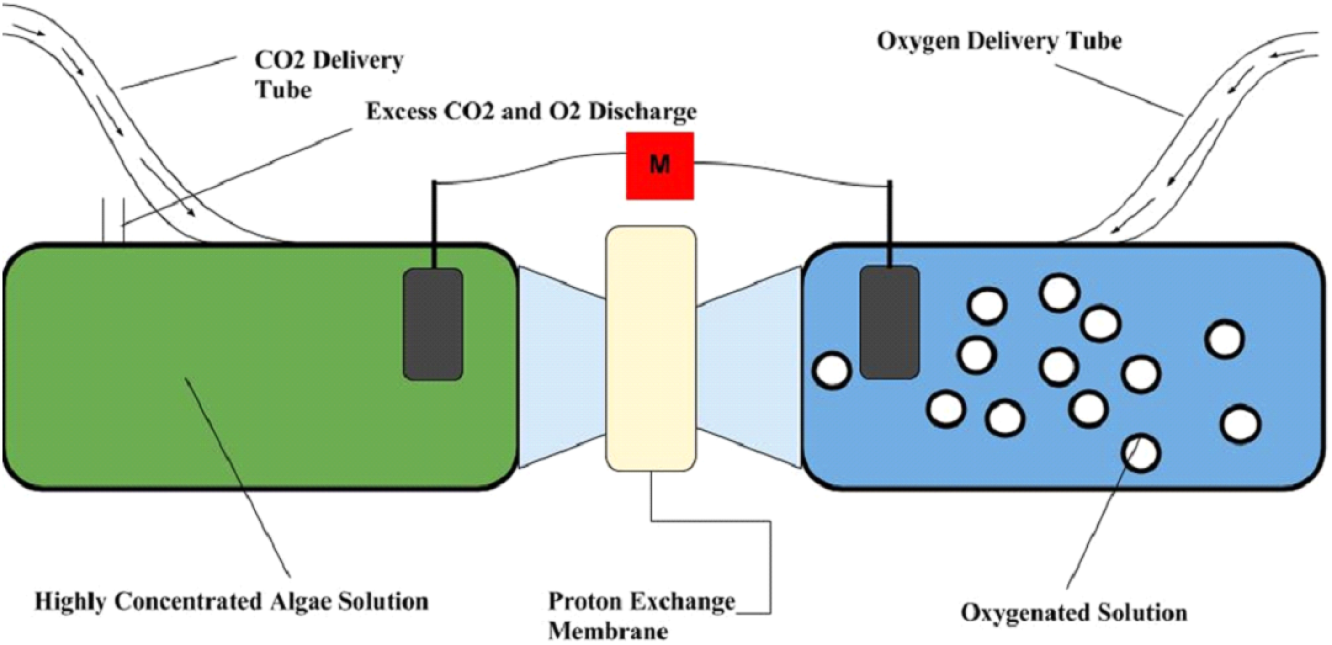
The 2D diagram further elaborates the technological specifications of the Microbial Fuel Cell System. The MFC system is first interlinked by a PEM (a sandwich structure) with an Anodic and Cathodic chamber. More importantly, the system includes two electrodes submerged into each chamber to gain electrical conduction.

**Figure 3:**
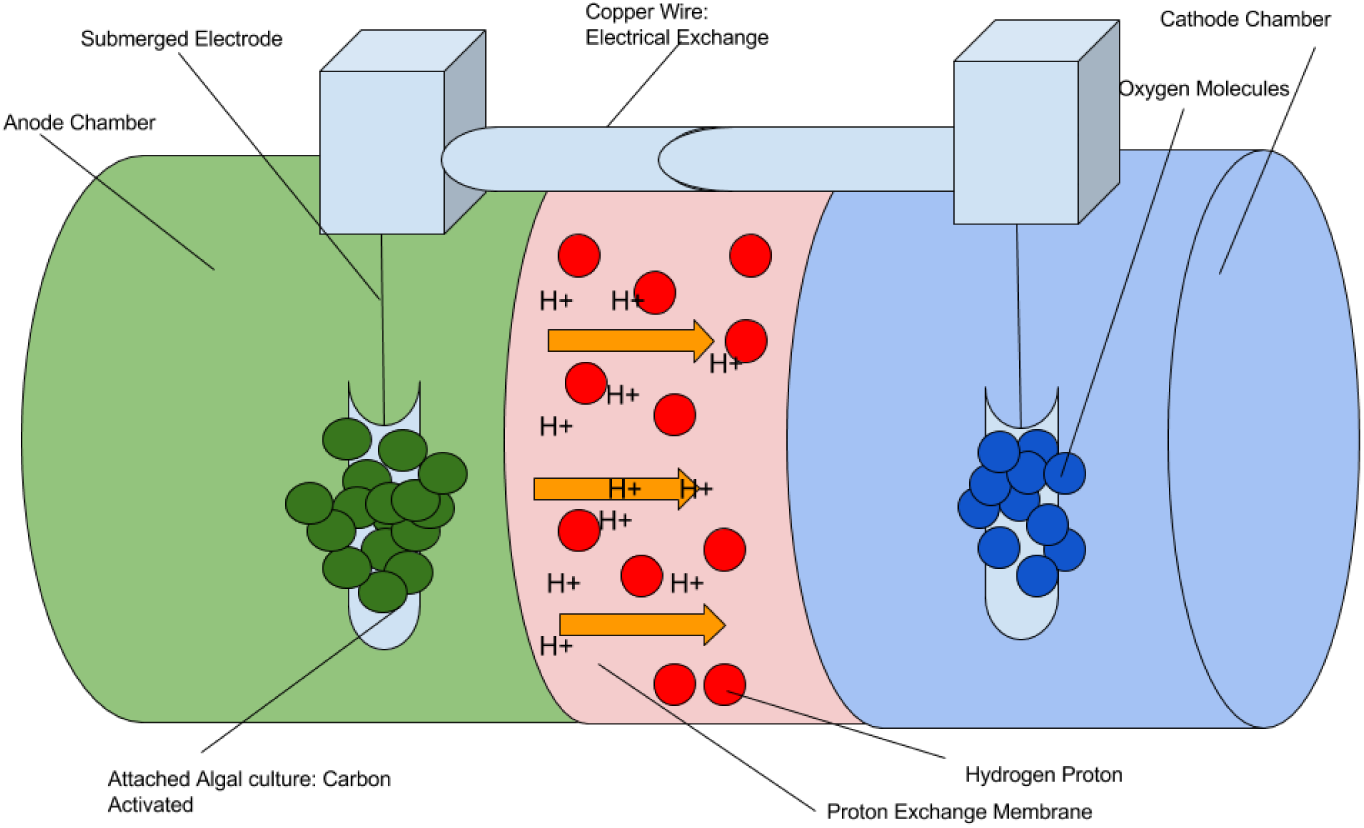
The compact Algal Microbial Fuel Cell is interlinked with a sandwiched Proton Exchange Membrane which exchanges Hydrogen protons between the Cathodic and Anodic chambers. The porosity of the PEM maximizes electrical output.

**Table 1.1:** Proton Exchange Membrane Mixture Protocol

|  |  |
| --- | --- |
| Mixed and Boiled Substance: Sodium Chloride | 75 (g) grams |
| Mixed and Boiled Substance: Water (H<sub>2</sub>O) | 200 (mL) milliliters |
| Mixed and Boiled Substance: Agar Media | 5 (g) grams |

> The mixture was later cooled until stable room temperature. The resulting mixture was then transferred into a petri dish. Finally, the petri dish was allowed to freeze in 0 degrees Fahrenheit or −17.78 degrees Celsius.
>
> The architecture of the Proton Exchange Membrane was unconventional in design, due to the technique of Hydrogen Proton Exchange. The substance reaction enabled maximal transfer proton transfer due to the medium’s high porosity. In other words, semipermeability was distributed across the membrane, rather than two singular location. Porosity was enabled with the reactants of the Agar medium. The function of the PEM is presented in the figure below:

### Organizing a Small Scale Unit of AlgaeBioreactors in Chlorella Vulgaris, Spirulina, and Nannochloropsis

Bioreactor design was most efficiently achieved with the traditional construction of the Microbial Fuel Cell system. A unified unit of varying Algal species (for the analysis of electrical output). The crucial components of a bioreactor are purposed to mimic the essential environmental conditions in a culturable Algal environment. The most significant Algae growth rate factors include nutrients and the photosynthetic access. Achieving an optimal ratio of sun exposure (3:1) (as identified in the parameters) in combination with nutrients levels was crucial (a Miracle Grow growth media). Additionally, a source of agitation was provided with a shaker platform as presented in the figure below:

**Figure 4:**
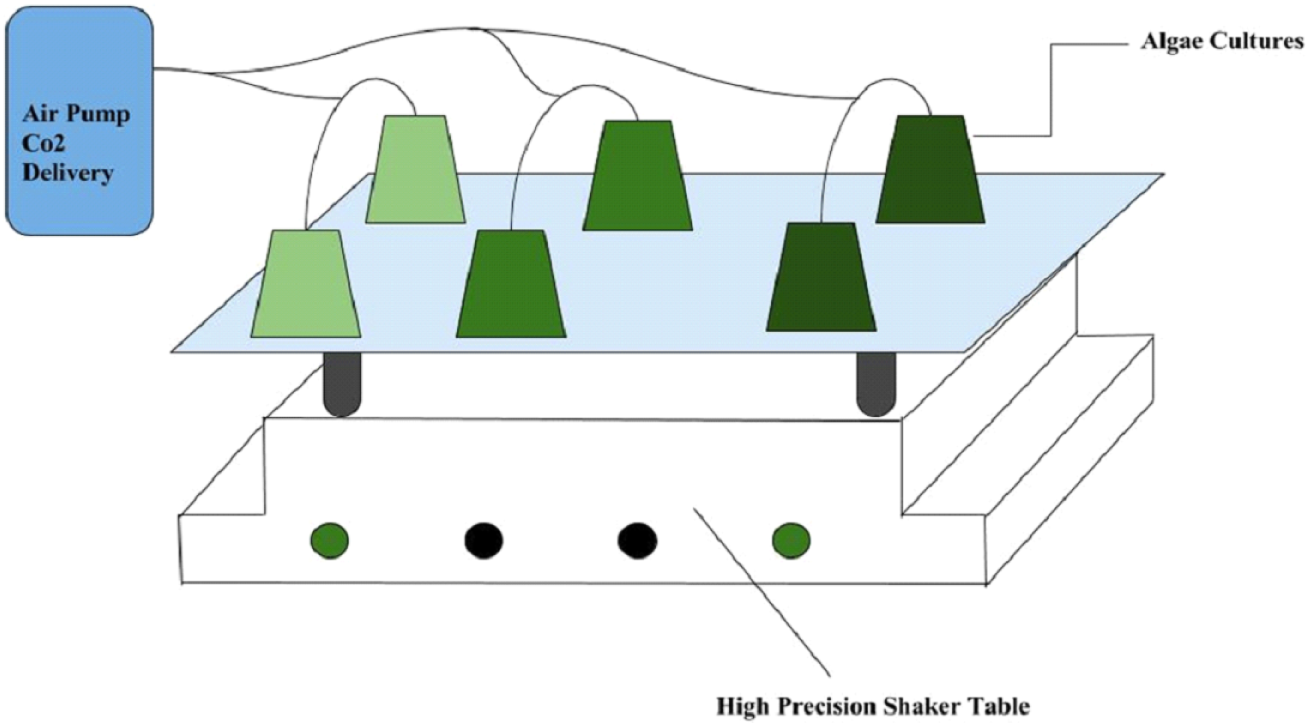
The diagram reveals an organized unit of Algal cultures in combination with the Microbial Fuel Cell system. A shaker platform was provided for continuous/systematic agitation. All parameters in each small-scale photo-bioreactor was regulated

### Algae Culturing Methods

Algae Culturing primarily consisted of the systematic maintenance of the chambers and the bioreactor system (the entire MFC system with its containment). Nearly 3.78 g/L of Miracle Gro (a generic nutrient media) was mixtured into the bioreactor system- or 0.003 g/mL. The bioreactor system primarily had two air pumps per microbial fuel cell system. This system also included close contact fluorescent lights to further maintain optimal growth conditions. Once the culture reduced towards 100 mL, it was replenished with 3.78 g/L of distilled water to replenish the bioreactor. Constant agitation was balanced through a laboratory shaker table at a setting of 2/10.

### Essential Materials

> Spectrophotometer: This will measure the absorptive frequencies of light, ensuring the growth rate of algae (this is a crucial part of the experimental procedure).

### Electrical output (mW) will be assessed and recorded by using a multimeter

> Potassium Hydroxide is used for lipid extraction, with nearly 1 ml of potassium hydroxide.

### Quantification of Variables: Electrical Output, Lipid Yields, and Cell Concentration

The variety of parameters and dependent variables obtained during experimentation included the resulting oil production, electrical output, and culture concentrations. Several instruments were utilized to precisely analyze each variable in relation to algae development. Cell concentration was primarily sampled to understand media and culture densities. In other words, algal density and absorption rates were recorded. A spectrophotometer was used to compare and acquire cell concentration. Higher algal culture absorption rates in (A) defined as highly dense cultures. Most importantly, electrical output was, simply, analyzed through a multimeter in mW electrical-energy. The multimeter was attached to a low-resistance copper wire which was coiled around a graphite stick submerged into the cathodic and anodic chambers of the MFC. Additionally, lipid yields were assessed to better understand the capabilities of oil production (in mL).

### Independent Variables Tested

An Algae-based microbial fuel cell will be compared in different culture species. More specifically, Nannochloropsis, Chlorella Vulgaris, and Spirulina will be investigated as an alternative biological energy source. Controlled Variables: The results obtained will than be compared with a generic bacterial-MFC system. The results, in terms of large scale energy production, will be analyzed, in relation to traditional fossil fuels and oil production.

### Constant Parameters

Throughout experimentation, parameters such as pH, salinity, lux levels (light intensity), temperature, and cell concentration were maintained. The containment was maintained by replenishing and releveling water levels to maintain constant cell concentrations. In addition, stabilizing pH, salinity, light intensity, and temperature was significant to ensure genuine results.

### Preliminary Conjectures

The hypotheses of the experimental trials is most associated with the scalability of Algae in combination with traditional MFC systems. Furthermore, the project explores the effectiveness of Algal culture as a sustainable substitute in small-large scale energy needs. The magnitude of a photosynthetic MFC is investigated by considering the potential electrical output and effectiveness of each Algal culture. Additionally, assessing the current provided understanding of the physical and biomolecular behaviors of each Algal genus is necessary (Chlorella Vulgaris, Spirulina, and Nannochloropsis) to hypothesize the magnitude of energy efficiency. As noted in the literature review, preliminary research indicates that Algal cultures cultivate at significantly distinct rates, in terms of cell concentration and reproductivity. Algae cultures, such as Spirulina, Chlorella Vulgaris, and Nannochloropsis were the prime candidates for MFC experimentation due to their notably high culture densities. Algal cells have high frequencies of cellular respiration and photosynthesis that are critical in MFC efficiency. The respiratory processes of an algae bioreactor enable high density algal growth, which occurs exponentially. Although an algae growth curve is S-selected, exponential growth occurred most commonly when approaching the stationary phase. Since algae cultures mimic similar exponential growth curves to that of aerobic and anaerobic bacteria, the following conjectures were formulated:

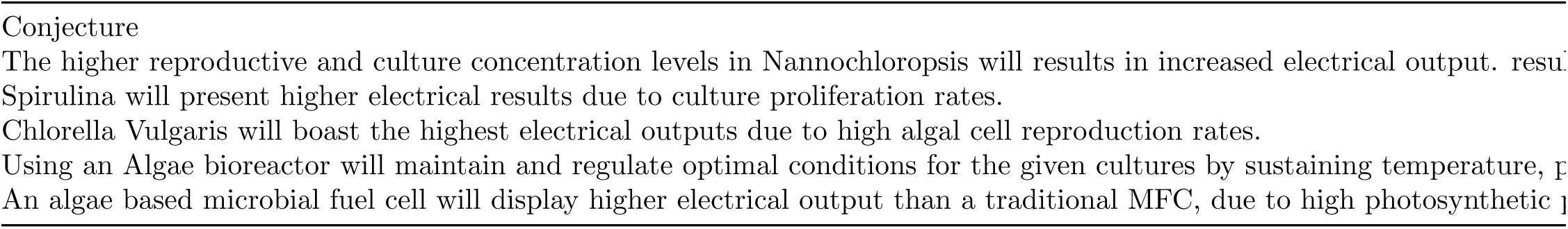

### Safety Risks and Factors of Experimentation

The most critical substance during the trials was Potassium Hydroxide (KOH) which was used for oil flocculation/extraction. Precautions, such as protective eyewear, gloves, and clothing were taken before this procedure. There are, however, are non-harmful methods of oil extraction such as compression. For the time being, Potassium Hydroxide was the most reachable method. This project does contain any other risk factors or safety concerns due to the materials included. The most concerning, however, is the algal culture and media. The algal culture was thoroughly maintained in a sterile environment and personal precautions were also taken-sterile protective gloves, eyewear, and clothing. By doing this we ensured personal safety and a controlled environment. The consumption of material and substances throughout the experimentation was prohibited.

## ASSESSMENT OF ALGAL CULTURE AND ELECTRICAL OUTPUT

After 50 mL of highly concentrated algae culture was obtained, it was immediately transferred to a 500 mL nutrient rich bioreactor medium composed of an initial 2 mg of All-Purpose growth media (1 mg/250ml). Cell concentration, pH, aquatic temperature, salinity, sterilization, and lux levels were assessed. Cell concentrations were obtained with a spectrophotometer. After two days of cultivation, the algae was transferred to a 450 mL MFC anode chamber. Electrical testing was executed with a multimeter. After all conditions and parameters were assessed in the anode and cathode chambers, cell concentration was maintained at a stationary phase in each algal culture bioreactor. Photosynthetic absorption levels served as the magnitude and benchmark of electrical production. The indirect correlation between absorption levels and electrical output were further investigated as a proportional increase from the stationary phase. Due to the proliferative nature of the stationary development, optimal electrical output (mW) were highly expected. The following photometric absorbance rates presented the most efficiency:

| Algae Species | Stationary Absorption Level |
| --- | --- |
| Chlorella Vulgaris | 123 (A) |
| Nannochloropsis | 286 (A) |
| Spirulina | 142 (A) |

### Data Analysis: Algae Cultivation

Throughout the analysis of data, there was an increasingly positive correlation between algae growth and electrical output until the stationary phase. However, additional experimental trials indicate that exceeding the stationary phase (death phase) will lower electrical output. Initially, electrical output was meager, however, nearing the stationary phase, mW increased drastically.

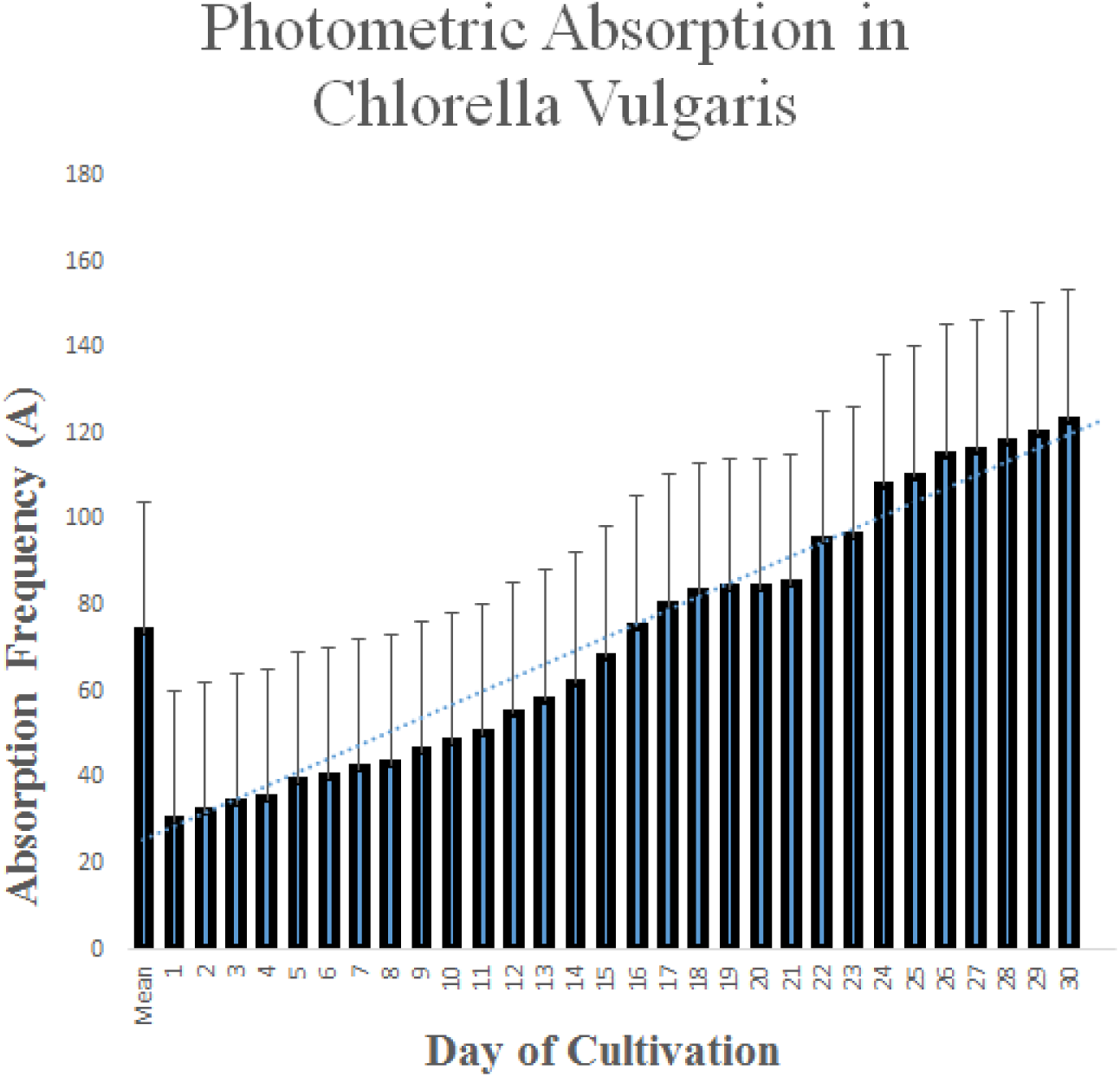

### Electrical Outputs in Individual Cultures

Peak Algal electrical outputs primarily occurred at the stationary phases of Algae development. In addition, the peak stationary absorption for Chlorella Vulgaris was 123 (A), Nannochloropsis 286 (A), and 142 (A) for Spirulina. Nannochloropsis presented the highest electrical output of 35 mW. Chlorella resulted in 30.2 mW, whereas Spirulina resulted in 31 mW. However, Spirulina experienced the most drastic increases in electrical outputs when approaching the stationary

**Figure 5:**
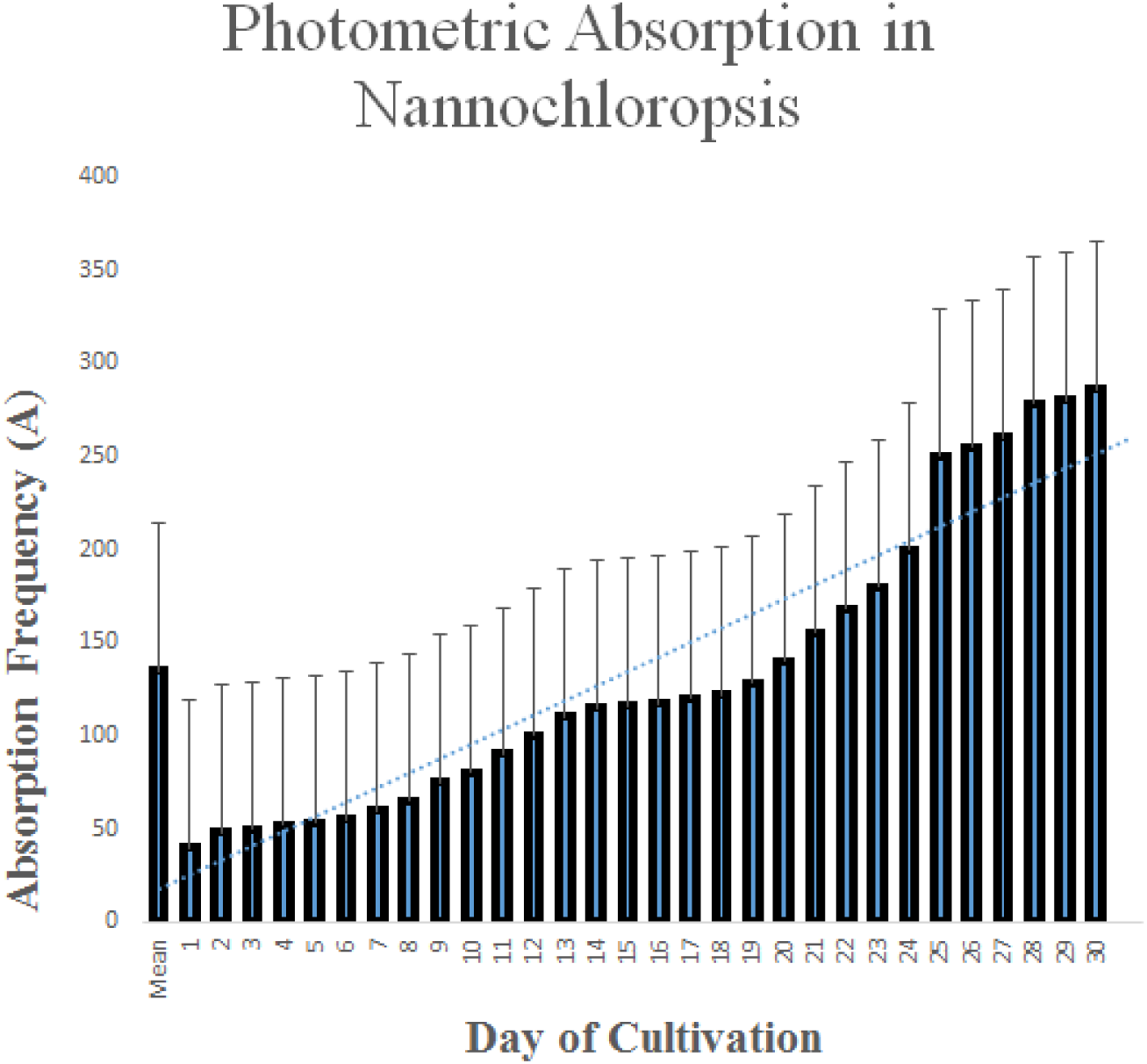
Physical data analysis of Algal development (in individual MFC systems) presented positive correlations between both Milliwattage and proliferation in the stationary phase. Due to high density cell densities in both Chlorella Vulgaris and Nannochloropsis, heightened metabolic rates were expected to satisfy the energy producing PEM. Chlorella Vulgaris peaked absorption levels at 123 (A), whereas, Nannochloropsis boasted a superior 286 (A).

**Figure 6:**
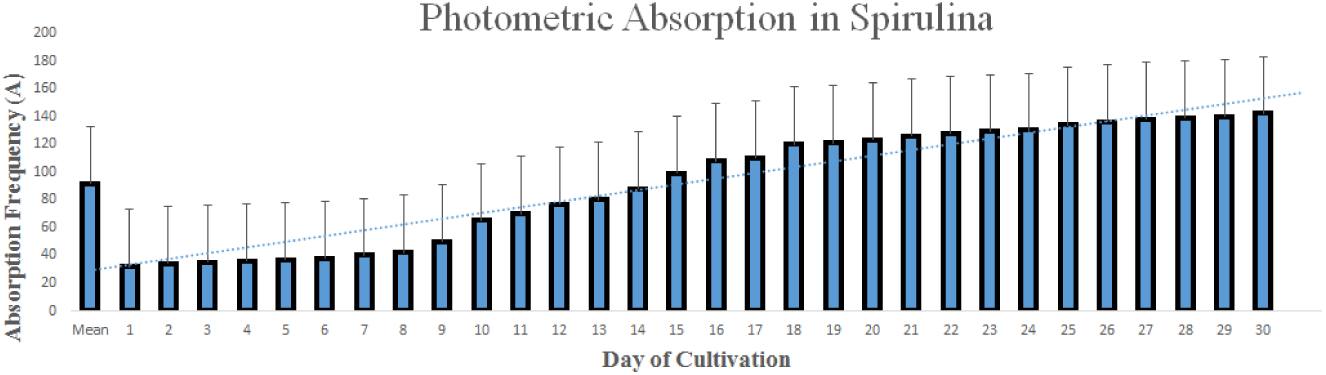
Spirulina development rates experienced less variation during the stationary phase. Although the stationary phase presents the highest source of physical density, Spirulina was nearly static and regulatory from days 18-30. However, in cultures such as Chlorella Vulgaris and Nannochloropsis, exponential growth was dispersed across the spectrum of the stationary phase. However, as noted with the previous cultures, proportional correlations were drawn between heightening electrical levels (mW) and absorption (A). Spirulina peaked a magnitude of 142 (A)

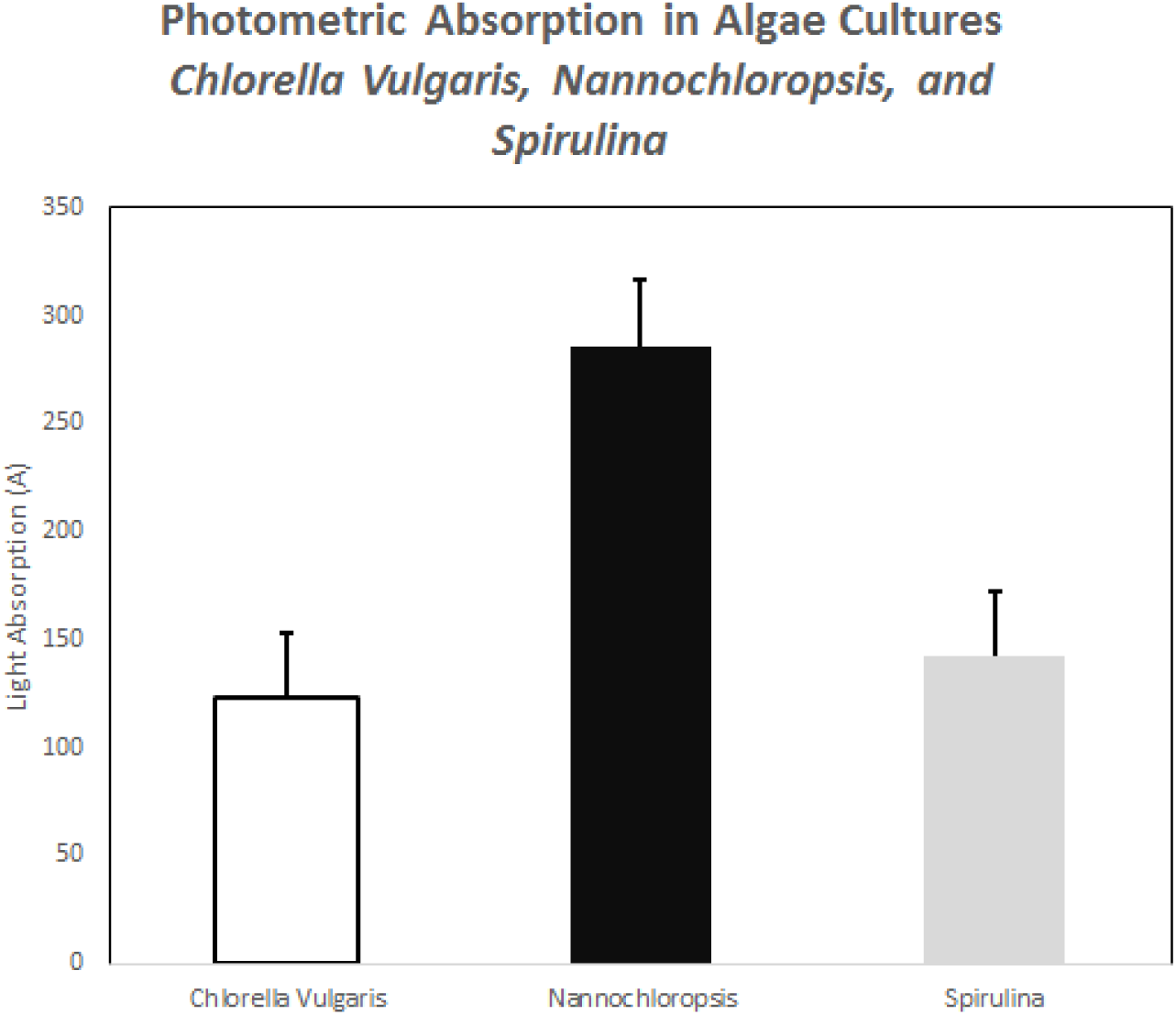

## SUPPLEMENTARY DATA

### Conclusions and Discussion

In this study, a novel algae-based photosynthetic MFC was evaluated in terms of electrical output, conditions, and growth. Within the electrical output, the nannochloropsis photometric absorption rate was at 286 (A) and had the highest electrical output of 35 mW. Additionally, the nannochloropsis was capable of proliferating during long periods of times. However, Spirulina (31 mW) and Chlorella (30.2mW) absorption rates proved to be less effective. It is likely that the diffusional limitation in the species resulted in an inefficiency of the absorption process. This preliminary study suggests that it is possible to utilize an algae-based microbial fuel cell to create energy efficient production through the photosynthetic nature of the fuel cell. Furthermore, the cell would reduce excess carbon dioxide while simultaneously producing energy. General Data Analysis: Electrical Output and Algae Development. During the investigation of data, it is concluded that Nannochloropsis is a likely candidate in bio-electric production. Most notably, there was an obvious correlation between both Algae development (photometric absorption) and electrical output mW. The positive relationship between electrical output (mW) and the magnitude of absorption (A) strongly suggests that higher algae concentrations were proportional with optimal electrical output when approaching the stationary phase. Due to higher metabolic processes in algal cultures, higher electrical results (mW) were anticipated.

### Future Directions and Prospects

The practical use of an MFC in a large scale setting is to be further investigated. An Algae MFC proves to be economically, environmentally, and technologically efficient. The future of this study may be used in multiple worldwide settings:

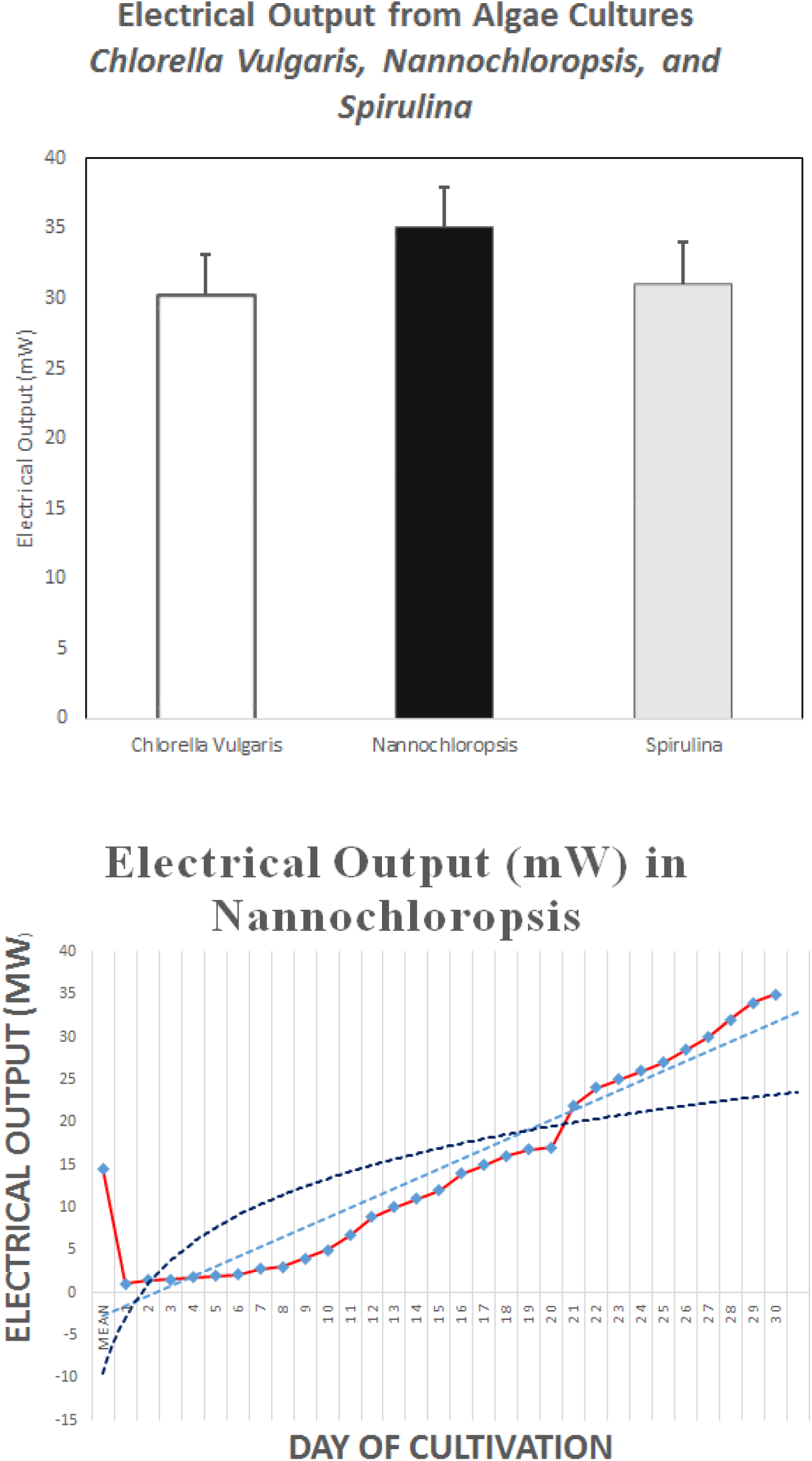

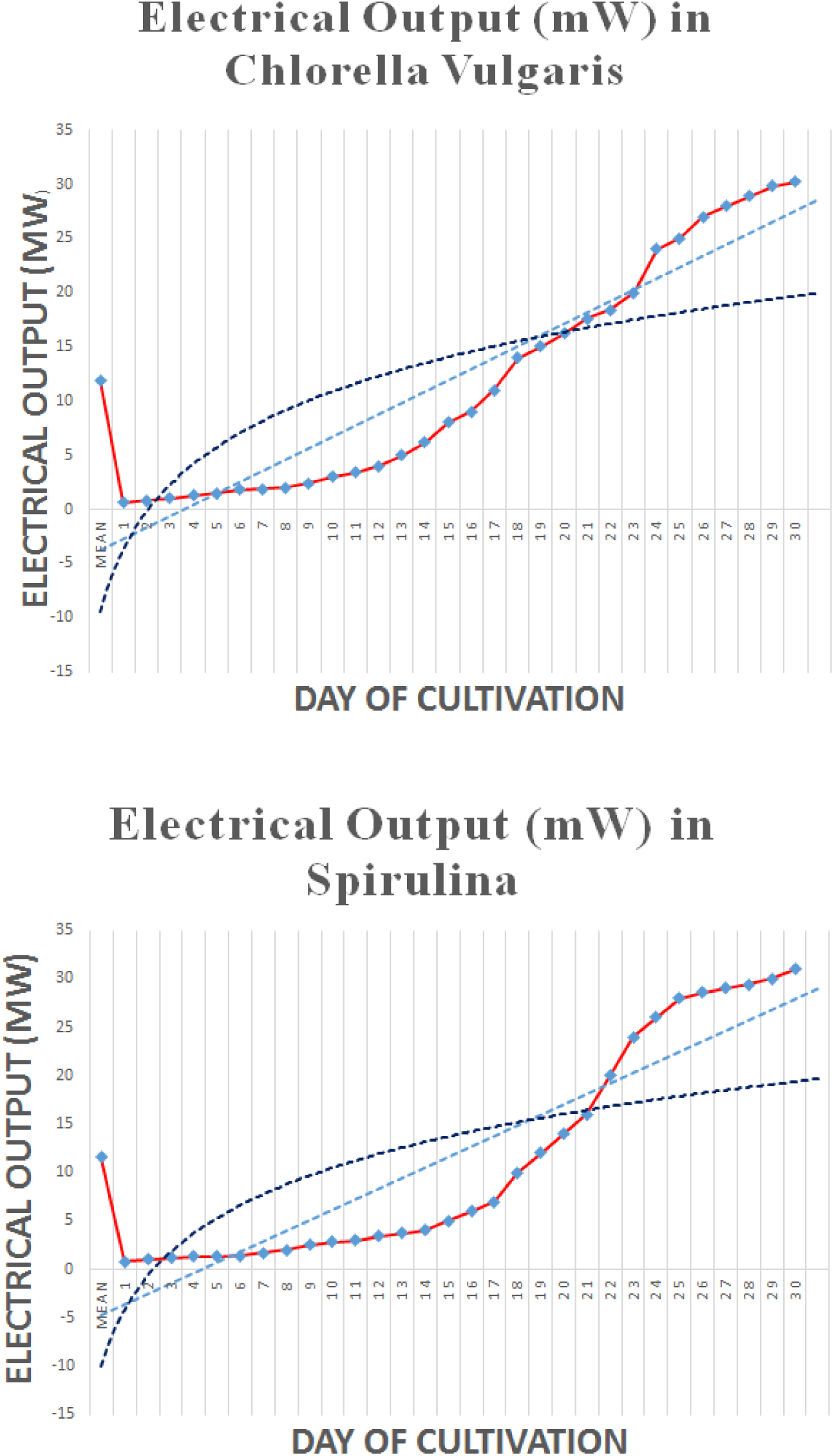

Developed Nations (e.g. United States and the European industry): Large-scale energy production as a substitute for fossil fuels and oil production. The MFC will immediately minimize greenhouse gases while producing biofuel and electrical energy.

**Figure 7:**
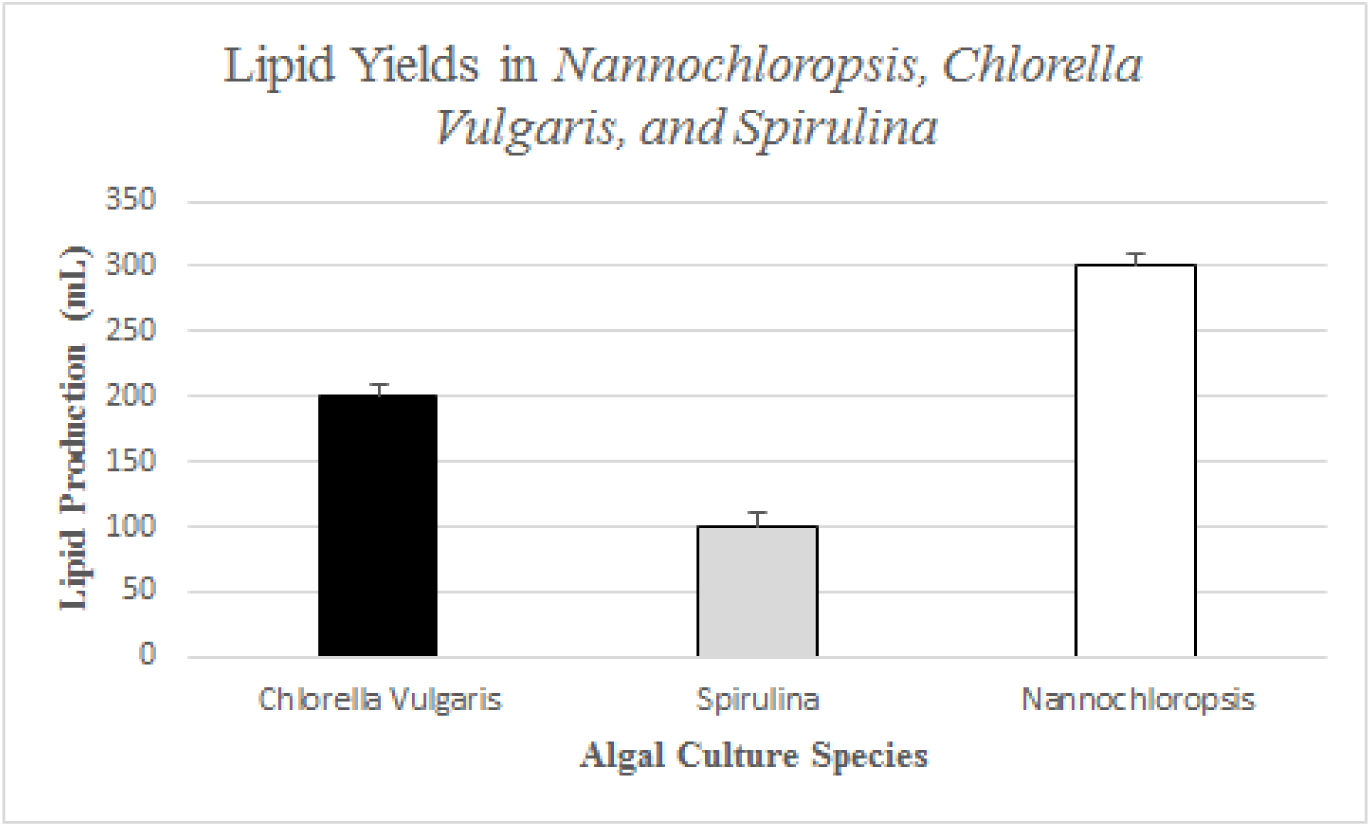
Although not the purpose of experimentation, Lipid Yields in varying algal cultures were assessed to acknowledge biofuel efficiency. The data presented indicates that nearly 200 mL of purified-natural oils were obtained from Chlorella Vulgaris, 100 mL in Spirulina, and 300 mL in a Nannochloropsis containment. Thus, the algal culture was able to simultaneously secrete oil yields while maintaining optimal electrical levels.

### Developing Countries (e.g. countries in Africa, Asia, and Latin America)

Potential Large scale electrical and biofuel production. May be used by individuals, facilities, and companies.

Large oceanic cultures can be utilized for high-scale energy production. More importantly, the scalability and size compatibility of a combined photobioreactor and microbial fuel cell is to be further sample. In addition, investigating the idea of small-scale interlinked Algal solar cells with maximized adjustability is crucial. Constructing a more adjustable unit of Microbial Fuel Cells may advantage easy scalability, attachment, and configuration (e.g. reorganization and repair of energy units after a storm or natural disaster).

